# Fasting reduces inhibitory and attentional control of food-related cues

**DOI:** 10.1101/2021.04.26.441416

**Authors:** M Ballestero-Arnau, B Rodríguez-Herreros, N Nuño-Bermúdez, T. Cunillera

**Affiliations:** Department of Cognition, Development and Educational Psychology. Faculty of Psychology, University of Barcelona; Pg. Vall d’Hebron, 171; 08035, Barcelona, Spain; Centre Cantonal Autisme, Centre Hospitalier Universitaire Vaudois and University of Lausanne, Rue du Bugnon 46; 1011, Lausanne, Switzerland; Universitat Oberta de Catalunya; Av. del Tibidabo, 39, 08035, Barcelona, Spain

## Abstract

The metabolic and cognitive systems interact to create the motivational drive that occasionally leads to disrupted consummatory eating behaviors. In this study, we investigated whether stimulus-specific alterations of the inhibitory function are present following a period of food deprivation. Twenty-six participants with normal weight performed the Stop Signal Task (SST) and the Go/No-go (GNG) task to measure response inhibition to food images containing high or low caloric content after following –or not- a 12-hour fasting period. Response inhibition performance in the SST did not exhibit significant differences when considering neither fasting, stimulus type nor food caloric content. We instead found a higher percentage of commission errors in the No-go trials of the GNG task in the fasting session, and specially with high-caloric food items. In contrast, the accuracy in the Go trials was similar between conditions. A mixed logistic regression model confirmed the remarkable impact of fasting on the performance of response inhibition. Overall, our findings support an interpretation of the motivational drive to eat strongly associated with aspects of the inhibitory function underlying high attentional control, rather than to a proper response inhibition per se.

## Introduction

Health systems in modern Western societies are already facing the direct consequences of normalizing the easy and frequent access to calorically dense and palatable foods (Swinburn et al., 2011; WHO, 2016). The ability to adjust behaviors such as controlling the amount of palatable food that one can eat –or abstain from eating it–, requires a well-established self-regulation in the cognitive system. Self-regulation can be understood as a form of inhibitory control (Baumeister, 2014), and considered as part of the brain executive functions (Miyake et al., 2000). In this vein, a deficient inhibitory control –and a poor self-regulation- are described in the development of impulsive control disorders (Groman, James, & Jentsch, 2009). The relationship between altered response inhibition processes in the brain and the presence of maladaptive eating behaviors has been widely established (Boisseau et al., 2012; Lavagnino, Arnone, Cao, Soares, & Selvaraj, 2016; Wu, Hartmann, Skunde, Herzog, & Friederich, 2013). Nevertheless, the specific cognitive mechanisms associated with halting food craving, especially under food deprivation conditions in which food cues turn to be a salient stimulus, are still poorly understood. In this study, we outlined how fasting modulates the inhibitory performance when coping with the desire to eat.

From an evolutionary perspective, complex neural mechanisms were shaped to react to immediate deficits in energy balance and, as a result, they were preserved to guarantee subsistence (Zeltser, Seeley, & Tschöp, 2012). However, the control of food intake - when accompanied by low energy consumption- was mostly neglected. Seminal studies on the mechanisms underlying the control of energy balance highlighted the debate between *metabolism* and *cognition*, and attempted to establish the role of each of these systems in controlling the motivational drive to eat. This unresolved question elicited a vast corpus of knowledge, revealing a powerful, distributed and redundant neural system in charge (Berthoud, 2002; Zheng & Berthoud, 2008). Cognition and metabolism interact to create the motivational drive that leads to consummatory eating behaviors (Zheng & Berthoud, 2008). More specifically, a metabolic need is translated into a behavioral action throughout the effect of hormones, transmitters and metabolic signals, leading to changes in neural excitability in corticolimbic structures (Berthoud, 2002).

It is commonly believed that we should avoid shopping for food while starving, or we could likely end up purchasing more items of higher caloric content than we had initially planned. Indeed, this altered food purchasing behavior induced by food deprivation has been consistently observed (Mela, Aaron, & Gatenby, 1996; Nederkoorn, Guerrieri, Havermans, Roefs, & Jansen, 2009; Nisbett & Kanouse, 1969). Under conditions of a metabolic deficit, behaviors become more impulsive and consequently more vulnerable to failures in the control of excessive food intake (Oliva, Morys, Horstmann, Castiello, & Begliomini, 2019; Wonderlich, Connolly, & Stice, 2004). Accordingly, impulsivity has been associated with insufficient inhibitory control and thus with greater difficulties in overruling automatic behaviors evoked by food stimuli, which can turn into an inability to control food intake (Nederkoorn, Houben, Hofmann, Roefs, & Jansen, 2010). In other words, converging evidence suggests that dysfunctional inhibitory control might be at the root of overeating (Claes, Nederkoorn, Vandereycken, Guerrieri, & Vertommen, 2006; Guerrieri, Nederkoorn, Stankiewicz, et al., 2007; Nederkoorn, Smulders, Havermans, Roefs, & Jansen, 2006). Likewise, the role of the inhibitory function in response selection is at the core of formal decision-making models that aim to describe how we decide and select appropriate actions (Bogacz, Brown, Moehlis, Holmes, & Cohen, 2006). Specifically, Nederkoorn and colleagues (Nederkoorn et al., 2009) showed the existence of a relationship between hunger and inhibitory function that partially explained patterns of excessive food consumption, particularly in dieters categorized as impulsive. Those participants were found to eat significantly more and to purchase more caloric food than controls, but importantly, only when feeling hungry.

Some studies have addressed the question of how hunger affects inhibitory function by employing the Go/No-go (GNG) task together with an attentional task (Kollei et al., 2018; Loeber, Grosshans, Herpertz, Kiefer, & Herpertz, 2013). They found that hunger affects not only the inhibitory function related to food consumption but also attentional control, especially in obese participants (Kollei et al., 2018). Likewise, studies that investigated the influence of hunger on the attentional bias phenomenon have reported that hunger enhances attention allocation towards food-associated cues in obese (Castellanos et al., 2009; Nijs, Franken, & Muris, 2009) as well as in normal-weight participants (Forestell, Lau, Gyurovski, Dickter, & Haque, 2012; Mogg, Bradley, Hyare, & Lee, 1998).

It could seem obvious that caloric food becomes a more salient stimulus under food deprivation conditions, as a short amount of food, but with a high caloric content, would be enough to return to the metabolic homeostasis and, consequently, to reduce the motivational drive to eat. Using different cognitive tasks, previous research has shown an attentional bias for food over non-food objects (Ballestero-Arnau, Moreno-Sánchez, & Cunillera, 2021; Kirsten, Seib-Pfeifer, Koppehele-Gossel, & Gibbons, 2019; Neimeijer, de Jong, & Roefs, 2013). More interesting for the purpose of the current study, several reports have proved the existence of a larger attentional bias for high-calorie food than for low-calorie food items (Cunningham & Egeth, 2018; Lee & Lee, 2021; Van Dillen, Papies, & Hofmann, 2013), whereas in other studies, an attentional bias for high-caloric items has been reported exclusively in a population with obesity (Bongers et al., 2015; Werthmann et al., 2011). Interestingly, Cunningham and Egeth (2018) evidenced that consuming a small amount of high-caloric food just prior to the experiment reduced the attentional bias found for high-calorie food to the level of low-calorie food items, demonstrating the high malleability of goal-states regarding the motivational drive to eat. With respect to the inhibitory function, obese individuals have shown deficits in the inhibitory function when high-calorie food items are used in comparison to low-calorie items (Gerdan & Kurt, 2020). However, in another study conducted on average weight women who binge eat, the authors did not find a deficit of the inhibitory function specific to high-calorie food cues, although those participants were highly responsive in a post-task food consumption from snacking high-calorie items (Lyu, Zheng, Chen, & Jackson, 2017).

Based on these premises, the goal of the current study is to investigate whether the inhibitory function is affected by the need to reduce the desire to eat that occurs after a period of fasting and to explore if this effect is specific of high valuable food items (high-calorie food) or instead it can be modulated by any kind of food-related cues. For this purpose, we designed an experiment in which participants performed two inhibitory tasks, the GNG and the Stop Signal Task (SST) (Huster, Enriquez-Geppert, Lavallee, Falkenstein, & Herrmann, 2013), responding to food or to nonfood images in two separate sessions: one following their normal eating habits and the other after a fasting period of approximately 12 hours. Furthermore, we manipulated the caloric content of food images in separate blocks to investigate how perceived calories is related to a possible decline of the inhibitory function. We hypothesized that if the motivational drive to eat affects inhibitory control related to food cues or exclusively to highly valuable food cues, it should be reflected in a poorer performance of the inhibitory function in both tasks following the fasting period but only when responses to food or high-calorie food items are the ones demanded to be inhibited.

## Method

### Participants

Thirty-three healthy volunteers participated in the experiment. None of them had a history of neurological deficits or eating disorders. All of them provided informed consent as approved by the local ethics committee prior to their participation in each experimental session. Data from 7 participants were discarded because the probability of responding on a stop trial [p(response|stop-signal)] was lower than 0.25 or higher than 0.75 (Verbruggen et al., 2019) for one or more conditions in the SST. Thus, the final sample was composed of 26 participants (24 females; *M* age = 20.8 ± 2.3 years; range: 18-26) with a normal body mass index (mean BMI = 22.6 ± 3.1). All participants were paid after completing the experiment.

A power analysis using G*Power (Erdfelder, Faul, Buchner, & Lang, 2009) indicated that, following the results obtained in previous studies that used a similar inhibitory task (Ganor-Moscovitz, Weinbach, Canetti, & Kalanthroff, 2018; Weinbach, Lock, & Bohon, 2020), a sample of 19 participants was sufficient to assess the effect of three within-subjects variables (with a power > 80% and a-priori alpha set at p =.05), using an effect size estimate of 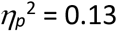.

### Stimulus

Three sets of 20 images were selected for the study from the foodcast research image database (FRIDa) (Foroni, Pergola, Argiris, & Rumiati, 2013). Image sets consisted of familiar pictures of high-calorie food (320.1 ± 124.3 kcal/100 g), low-calorie food (88.1 ± 96.9 Kcal/100 g), and office supplies. Both sets of food images contained natural and prepared food. All three sets of images were comparable in terms of size, spatial frequency, brightness, familiarity, typicality, and ambiguity. However, compared with the office supply images, the food images yielded significantly higher values for valence [food = 68.15; office supply = 61.91; *t*(19) = 1.985; *p =* 0.054] and arousal [food = 46.99; office supply = 28.31; *t*(19) = 4.48; *p <* 0.001]. A sound (22050 Hz, 200 msec, 5 msec ramp on and off) was used as a stop signal. Another two sets of 20 images, food, and kitchen supplies, were selected following the same procedure, to be used in the practice blocks.

### Inhibitory paradigms

We implemented the two most used tasks for studying the inhibitory function, the GNG task and the SST, to observe, at the within-subject level, how hunger affects the inhibitory process. The same stimulus, number of trials, stimulus presentation duration, and response pattern were used in both tasks. The images of food and office supplies served as stimuli in both tasks and were presented on a screen (white background), subtending 8.9° of the visual angle.

In the GNG task, one stimulus was presented 2.4 cm on the left or right side of a permanent central fixation cross at a time, while in the SST, stimuli were presented, one at a time, at the center of the screen, replacing the central fixation cross. In each trial and for both tasks, the stimulus was presented for 500 msec, and the stimulus onset asynchrony (SOA) was fixed across trials and randomized between participants and sessions separately for the Go and No-go trials in the GNG task and for the Go and Go + Stop trials in the SST (range: 1100-1500 msec; mean = 1315 msec ± 118).

The GNG task consisted of 8 blocks of 100 trials, in which 75% of the trials corresponded to Go responses and the other 25% corresponded to No-go responses. In 4 consecutive blocks, food images were assigned as Go and office supplies images as No-go, and vice versa for the other 4 blocks. In two consecutive blocks, food images corresponded to high-calorie items, whereas for the other two blocks low-calorie food images were used. The SST consisted of 4 blocks of 200 trials, in which 75% of the trials corresponded to Go responses and the other 25% corresponded to Go + Stop responses. The stop signal was presented following a variable delay after stimulus onset. The stop-signal delay (SSD) was initially set to 250 msec at the beginning of each set of 2 consecutive blocks with different category-response assignments and was adjusted separately for food and nonfood conditions. The SSD was adapted to each participant’s behavior by means of a staircase-tracking algorithm (Logan, Schachar, & Tannock, 1997). Thus, the SSD was increased or decreased by 25 msec after successful or unsuccessful response inhibition, respectively. This dynamic tracking procedure guarantees an overall p(response|stopsignal) of 0.5. High-calorie and low-calorie food images were presented in different blocks, and block order and category-response assignment were counterbalanced across participants and sessions in both tasks.

Participants were required to respond using left- and right-hand responses with the corresponding index finger in both tasks. Importantly, the response hands were equally frequent and pseudorandomly distributed within each block of the experiment. Thus, for the GNG task, participants responded on the side where images in the category assigned as Go responses appeared and were asked to withhold responding to the other category of images (No-go condition). The category-response assignment was reversed after completing half of the experiments, and participants were informed at the time. Analogously, for the SST, the image category was assigned to right/left responses, and participants were required to classify images accordingly but to withdraw a response whenever they heard the stop-signal sound, presented through headphones. Again, the category-response pattern was reversed after completing half of the task and notifying participants.

Finally, the following constraints were introduced in both tasks: i) no more than three consecutive stimuli appeared on the same side (GNG), or the same image category (food or nonfood) was not presented in more than three consecutive trials (SST), ii) two consecutive No-go or Go + Stop trials never occurred, and iii) the same stimulus was never immediately repeated.

A practice block of 40 trials was presented prior to each task, with 50% of the trials corresponding to No-go or Go + Stop trials to guarantee a full proof and a complete understanding of the tasks at hand. In the SST, only two equally distributed SSD values, 100 and 500 msec, were implemented. The practice block for the GNG task was repeated when the error rate was larger than 30% or the average reaction time (RT) was longer than 700 msec, while for the SST, the practice block was repeated when the percentage of inhibitory errors was larger than 50%, the error in the Go trials was larger than 20%, or the average RT was slower than 700 msec.

### Procedure

Each subject participated in three sessions. In the first one, participants went through an anamnesis to rule out any psychological or eating disorder that could make undergoing an ~12 hour fasting period difficult. Weight and height data were collected from all participants. Finally, participants completed the EDI-2 questionnaire (Seibt, Häfner, & Deutsch, 2007).

The experimental sessions took place in the morning (~10.30 a.m.). The session order was counterbalanced across participants, and the intersession intervals were at least one week long. In the fasting session, participants were asked to refrain from consuming any solid or liquid food other than water after 10.30 p.m. of the previous day, while in the nonfasting session, participants were asked to follow their usual eating habits before participating in the experiment. Each task always began with the practice block. Short breaks of 15 sec were included in each task every hundred trials to allow participants to rest. Each experiment session lasted for ~55 min.

### Analytical measurements and statistical analyses

For each participant, the blood glucose concentration and hunger feeling were measured and subjected to analysis of variance and *t*-tests, respectively. The former measurement was accomplished using a glucometer (SD Biosensor Inc.), following a standardized safety protocol (“Infection Prevention during Blood Glucose Monitoring and Insulin Administration,” n.d.). Hunger feeling was assessed by asking participants to report their hunger level on a rating scale that corresponded to a 10 cm line drawn on a sheet of paper with marked intervals from 0 to 10 points, where 0 meant *not hungry at all* and 10 corresponded to *starving*. Blood glucose levels were measured twice, before and after completing the experiment in both experimental sessions, while the subjective rating was measured at the beginning of the experimental session.

Behavioral measurements of Accuracy on Go trial, reaction times (RT) on correct Go trials, and accuracy on Nogo and Stop trials were extracted from both inhibitory tasks, while RTs on unsuccessful stop trial and the computation of SSRTs were only obtained in the SST. The inhibitory function in the SST was measured using the SSRTs, which was estimated separately for food and nonfood conditions, as well as for high- and low-caloric content conditions. We used the integration method to calculate the SSRT (Verbruggen & Logan, 2009). We evaluated all the parameters of the participants’ performance in the SST and the GNG task in separated 2 × 2 × 2 repeated measure analysis of variance (ANOVA), with the within-subject factors *fasting* (hungry vs. satiated), *image category* (food vs. non-food), and *caloric content* (high-calorie vs. low-calorie). All data were log-transformed prior statistical analyses to normalize their distribution (Ratcliff, 1993).

Finally, the inhibitory performance in the GNG task was further analyzed by fitting the binary data (correct-error) from the No-go trials into a logistic regression model using the R-package lme4 (Bates, Mächler, Bolker, & Walker, 2015). The factors *fasting*, *image category* and *caloric content* were entered in the model as fixed effects, while the intercepts for subjects, items and SOA were entered as random effects. The overall fit of each effect was assessed using *p*-values obtained by the likelihood ratio test, comparing the model with the effect against the same model without the effect. The simplest model included *image category* as a fixed factor and by-items and by-SOAs as random intercepts. The random intercept by-subjects was incorporated into the model when *fasting* was added to the model.

## Results

The food deprivation period was effective modulating both hunger feeling [fasting session: 7.3 ± 2.2; nonfasting session: 3.7 ± 2.4; *t*(25) = 6.3; *p <* 0.001; *d =* 1.56] and blood glucose levels [fasting: *F*(1,25) = 15.2; *p =* 0.0006; 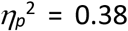; pre-post-experiment: *F*(1,25) = 8.8; *p =* 0.007; 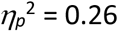; fasting x pre-post-experiment *p*-value = 0.8].

A summary of the performance in the SST is presented in Table 1. The results revealed that, on average and across conditions, accuracy on Go trials reached 87.5% (±6.4 S.D.). Participants’ accuracy on Go trials was similar between fasting and nonfasting sessions [*F*(1,25) = 3.0; *p =* 0.095; 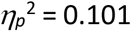]. Similarly, no differences were observed for *caloric content* [*F*(1,25) = 0.093; *p >* 0.7; 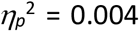], whereas we found a trend for higher accuracy responding to food than to non-food images [*F*(1,25) = 3.77; *p =* 0.064; 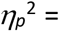 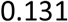]. No difference in RTs were observed for *fasting* [*F*(1,25) = 0.138; *p >* 0.7; 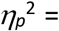 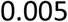], *image category* [*F*(1,25) = 0.742; *p >* 0.3; 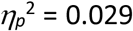], or *caloric content* [*F*(1,25 = 0.008; *p >* 0.9; 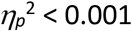; see Figure 1].

**Table 1.** Mean values and standard deviations for the *fasting* and *image category* conditions with *low-high-caloric* food in the in the Stop Signal Task (SST). The presented results correspond to the mean accuracy and reaction time (RT) in Go trials. The Stop Signal Reaction Time (SSRT) was calculated following the integration method.

| Accuracy (%) Go trials |  |  |  |  |  |  |  |  |
| --- | --- | --- | --- | --- | --- | --- | --- | --- |
|  | Fasting |  |  |  | Nonfasting |  |  |  |
|  | Food |  | Nonfood |  | Food |  | Nonfood |  |
|  | M | S.D. | M | S.D. | M | S.D. | M | S.D. |
| *High-calorie* | 89.5 | 6.8 | 87.9 | 14.3 | 86.3 | 8.1 | 85.8 | 9.2 |
| *Low-calorie* | 90.8 | 5.1 | 87.3 | 8.4 | 86.9 | 7.6 | 84.4 | 9.5 |

| RT (msec) Go trials |  |  |  |  |  |  |  |  |
| --- | --- | --- | --- | --- | --- | --- | --- | --- |
|  | Fasting |  |  |  | Nonfasting |  |  |  |
|  | Food |  | Nonfood |  | Food |  | Nonfood |  |
|  | M | S.D. | M | S.D. | M | S.D. | M | S.D. |
| *High-calorie* | 499 | 131.4 | 501 | 128.9 | 508 | 124.8 | 512 | 134.4 |
| *Low-calorie* | 504 | 132.7 | 510 | 134.1 | 501 | 105.0 | 502 | 114.3 |

| SSRT (msec) |  |  |  |  |  |  |  |  |
| --- | --- | --- | --- | --- | --- | --- | --- | --- |
|  | Fasting |  |  |  | Nonfasting |  |  |  |
|  | Food |  | Nonfood |  | Food |  | Nonfood |  |
|  | M | S.D. | M | S.D. | M | S.D. | M | S.D. |
| *High-calorie* | 202 | 47.0 | 209 | 50.7 | 210 | 71.6 | 211 | 70.0 |
| *Low-calorie* | 191 | 83.3 | 207 | 48.2 | 214 | 76.3 | 216 | 84.4 |

**Figure 1.**
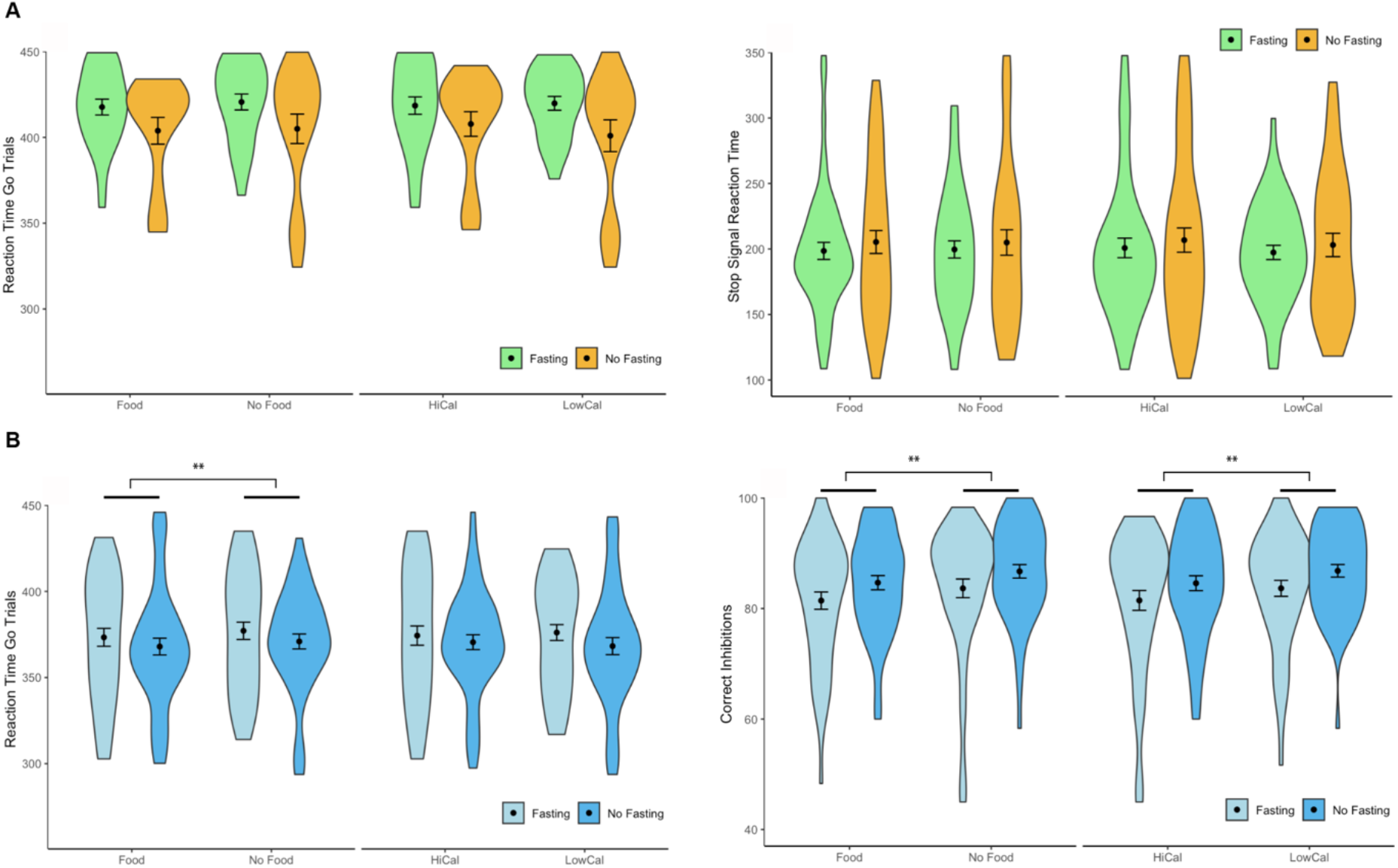
**A)** Results of the SST (RTs and SSRTs) and **B)** Results of GNG task (RTs and Nogo hits). All these results are presented separately for *fasting*, *image category*, and *caloric content*. The asterisk indicates statistically significant differences (** = *p*-value < 0.01).

The RTs in unsuccessful stop trials (462 msec) were faster than in correct Go trials [505 msec; *t*(25) = 8.26; *p <* 0.0001; *d =* 0.41]. This is in line with the statement of the horse race model about the context independence assumption, that serves to validate the computation of the SSRTs. In brief, this assumption states that the stop and go processes are unrelated, thus allowing to used go RT on no-stop-signal as an estimate of go RTs on stop-signal trials to predict the signal-respond RTs and then, to calculate the SSRT (Verbruggen & Logan, 2017). Likewise, we observed that Stop accuracies were close to 50% (mean percentage correct stop trials = 51.6 ± 6.7%), and that no difference in inhibitory performance was found across conditions, altogether indicating that the current version of the SST worked as expected and therefore, the inhibitory performance in the SST was examined. The results of the analyses of the SSRT revealed that fasting did not modulate participants’ inhibitory performance [*F*(1,25) = 1.072; *p >* 0.3; 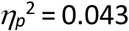; see Figure 1]. No effects of image category, caloric content or significant interactions were found for the SSRT (all *p*-values > 0.2).

The results of the GNG task are summarized in Table 2. Participants’ accuracies on Go trials, averaged across conditions, was 99.0% (± 1.3 S.D.). The accuracy in the Go trials was comparable in all conditions [*fasting*: *F*(1,25) = 0.473; *p >* 0.4; 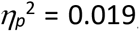; *image category*: *F*(1,25) = 0.197; *p =* 0.661; 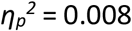; *caloric content*: *F*(1,25) = 1.564; *p >* 0.2; 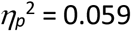]. None of the interactions reached statistical significance (all *p*-values > 0.4). RTs were similar in the fasting and nonfasting sessions [*F*(1,25) = 0.519; *p >* 0.4; 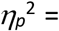 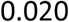] and for the high- and low-calorie conditions [*F*(1,25) = 0.008; *p >* 0.9; 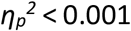]. However, RTs were faster for the food (373msec) than for the nonfood (381msec) condition [*F*(1,25) = 14.467; *p =* 0.001; 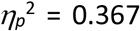]. Again, none of the interactions reached statistical significance (all *p*-values > 0.1). The influence of fasting in the inhibitory performance of Nogo trials was marginally significant [*F*(1,25) = 3.658; *p =* 0.067; 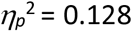], whereas a significant effect of *image category* [*F*(1,25) = 8.827; *p =* 0.006; 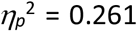] and *caloric content* [*F*(1,25) = 11.699; *p =* 0.002; 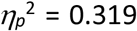] was found (see Figure1). The interaction *image category* x *caloric content* was also marginally significant [*F*(1,25) = 4.103; *p =* 0.054; 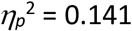].

**Table 2.**
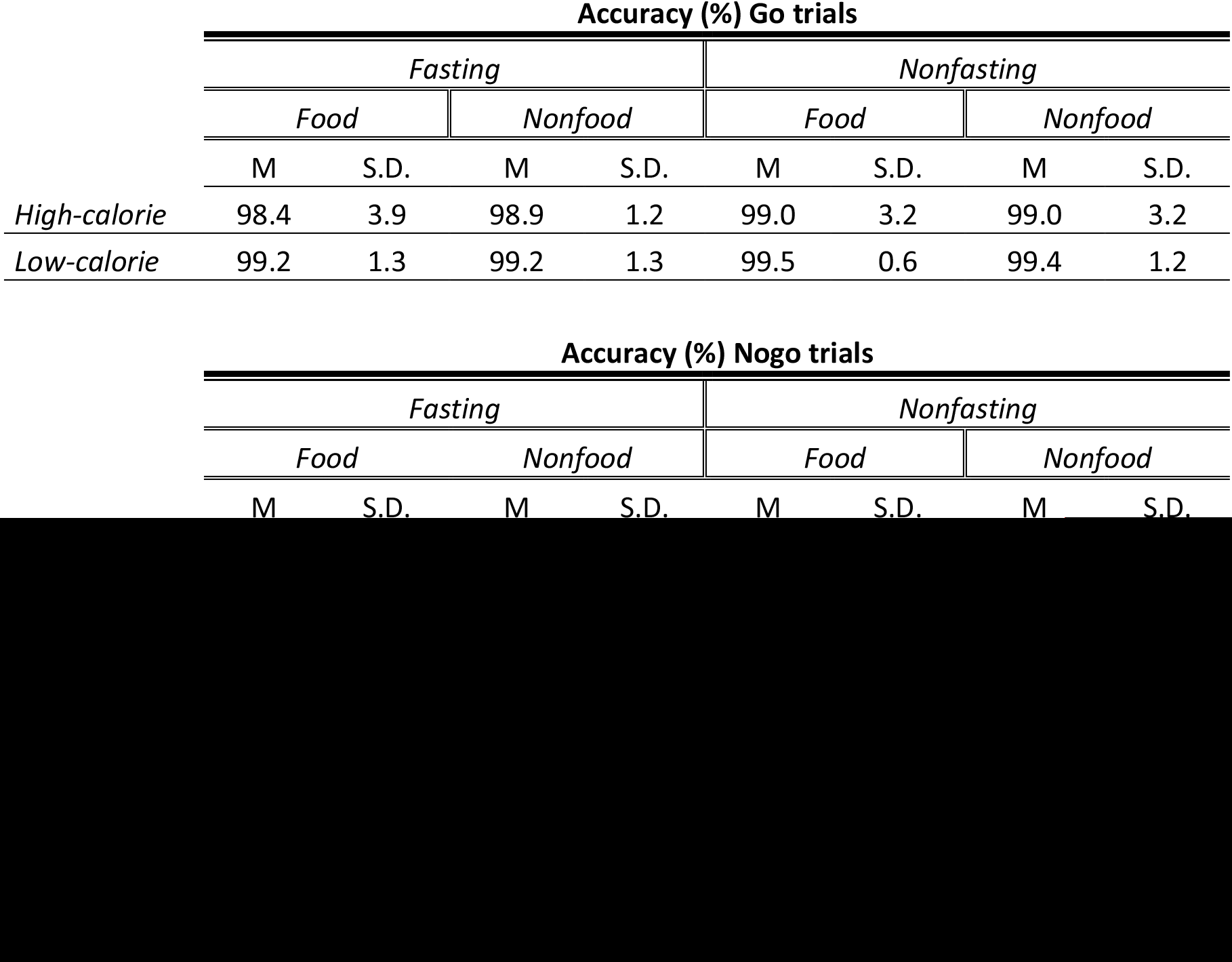
Mean values and standard deviations for the fasting and image category conditions with low- high-caloric food in the Go-Nogo (GNG) task. The presented results correspond to the mean accuracy and reaction time (RT) in Go trials. Mean accuracy for Nogo trials is also presented and correspond to the measurement of response inhibition in this task.

We further analyzed the inhibitory function in the GNG task using a logistical linear regression model. First, the results indicated that convergence was reached in all models constructed. The estimates of the full model are reported in Table 3. Crucially, the model in which image category and caloric content were assigned as fixed factors was compared with another model in which fasting was added as a fixed factor (see Figure 2). This comparison revealed an improvement in the model fit [*χ*^2^(2) = 105.7; *p <* 2e-16].

**Table 3.** Estimates and results of the three mixed logistic regression models fitting the accuracy of the response inhibition in the GNG task. Comparison of the AIC and BIC of the three models and of the results of the likelihood ratio test.

| Model 1<br>*image category* |  |  |  |  | Model 2<br>*Image cat. + caloric-content* |  |  |  | Model 3<br>*Image cat + caloric-cont. + Fasting* |  |  |  |
| --- | --- | --- | --- | --- | --- | --- | --- | --- | --- | --- | --- | --- |
| Random effects | *Variance* |  | *SD* |  | *Variance* |  | *SD* |  | *Variance* |  | *SD* |  |
| Item | 0.182 |  | 0.43 |  | 0.176 |  | 0.42 |  | 0.18 |  | 0.42 |  |
| ITI | 0.61 |  | 0.78 |  | 0.61 |  | 0.78 |  | 0.15 |  | 0.38 |  |
| Participant | - |  | - |  | - |  | - |  | 0.43 |  | 0.66 |  |
| Fixed effects | *Estimate* | *SE* | *z-value* | *p-value* | *Estimate* | *SE* | *z-value* | *p-value* | *Estimate* | *SE* | *t-value* | *p-value* |
| (Intercept) | 1.64 | 0.17 | 9.79 | <2e-16 | 1.59 | 0.17 | 9.36 | <2e-16 | 98.73 | 0.33 | 294.96 | < 2e-16 |
| Food/Nonfood | 0.27 | 0.13 | 2.11 | **0.035** | 0.27 | 0.13 | 2.14 | **0.032** | 0.27 | 0.13 | 2.14 | **0.033** |
| Caloric content | - | - | - | - | 0.09 | 0.07 | 1.39 | 0.17 | 0.09 | 0.07 | 1.39 | 0.17 |
| Fasting | - | - | - | - | - | - | - | - | 0.23 | 0.11 | 2.12 | **0.034** |

| Model Fit Criteria |  |  | LRT |  |  |
| --- | --- | --- | --- | --- | --- |
| MODEL | *AIC* | *BIC* | *Chi-sq* | *df* | *p-val* |
| Model 1 vs Model 2 | 10394.7 vs 10394.8 | 10424.5 vs 10432 | 1.898 | 1 | 0.17 |
| Model 1 vs Model 3 | 10394.7 vs **10293.3** | 10424.5 vs **10345.5** | 107.4 | 3 | **<2e-16** |
| Model 2 vs Model 3 | 10394.8 vs **10293.3** | 10432 vs **10345.5** | 105.7 | 2 | **<2e-16** |

**Figure 2.**
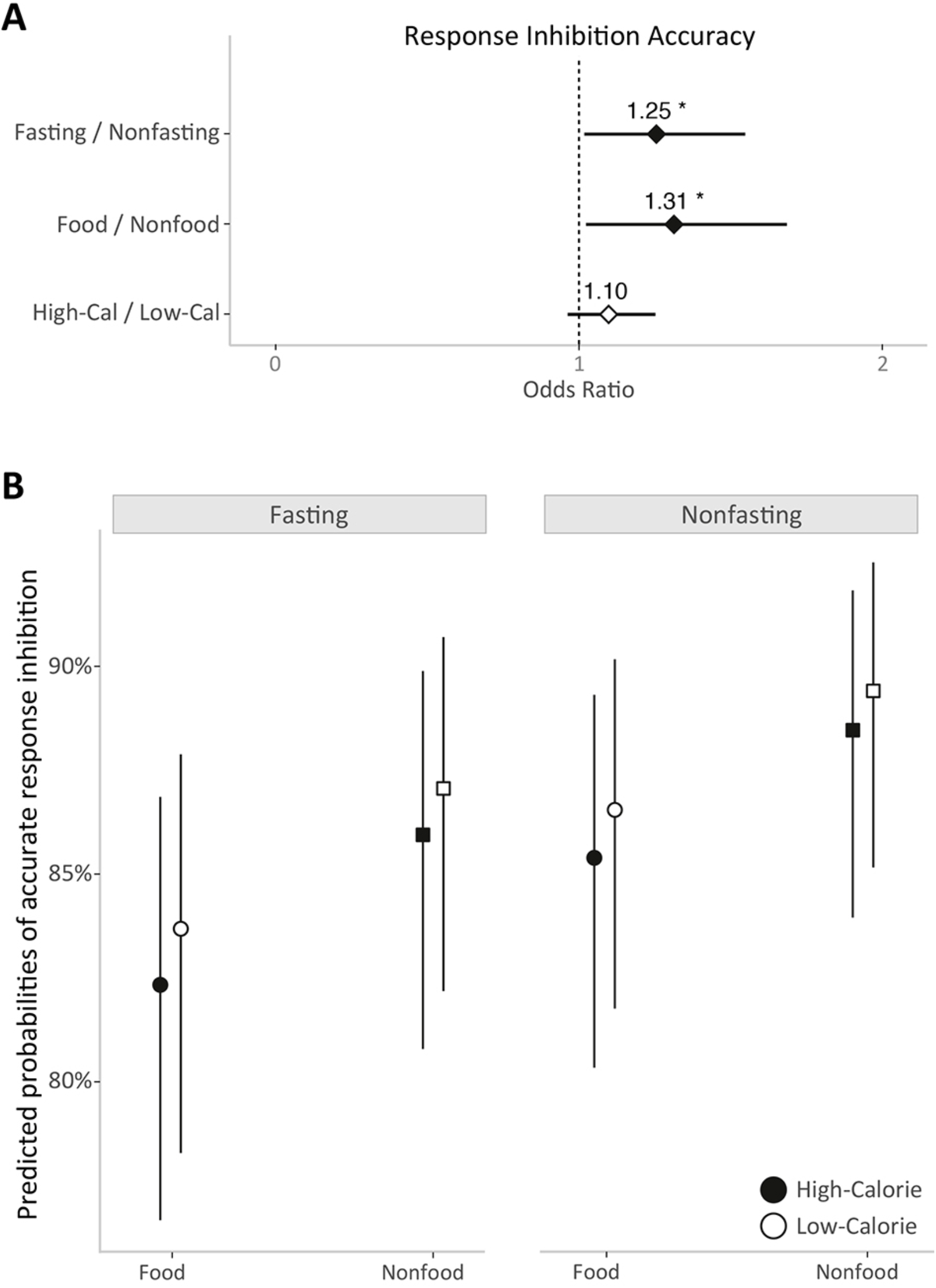
**A)** Forest plot of the odds ratios for all fixed factors (*image category*, *caloric content* and *fasting*) of the mixed effects logistic regression that fits the accuracy of response inhibition in the GNG task. Error bars represent 95% confidence intervals. The asterisk indicates statistically significant differences (* = *p*-value < 0.05). **B)** Predicted percentages of accurate response inhibition by the mixed effects logistic regression for each image category (food in circles, nonfood in squares) and caloric content (high-calorie in black, low-calorie in white) under fasting and nonfasting conditions.

## Discussion

In the present study, we evaluated the influence of inducing a motivational drive to eat on inhibitory performance when food and nonfood cues were presented, and especially when food images varied in their caloric content. To that end, we used the two most widely used paradigms to investigate the inhibitory function: the SST and the GNG task. Our results demonstrated that food deprivation has a general effect on the inhibitory performance, but it was only observed in the GNG task. Furthermore, we found that participants had significantly more difficulty withholding their response to food than to nonfood images, and this effect was larger for high-calorie than for low-calorie food items. Thus, our participants had no difficulty withdrawing responses regardless of the food deprivation state or the experimental conditions, as measured by the SSRT in the SST. These results suggest a behavioral effect related to the motivational drive to eat and linked to some specific aspects of what is considered the inhibitory function.

The inhibitory function has been described to require the interplay between signal detection, action selection, and action execution or suppression (Verbruggen, McLaren, & Chambers, 2014). Our results point to the fact that the motivational drive to eat may affect signal detection and action selection in a different way in the two inhibitory tasks, which was reflected in a detrimental inhibitory performance observed exclusively in the GNG task. Thus, although the inhibitory process engaged in both inhibitory tasks was driven by consistent stimulus response mapping, successful stop inhibition in the SST relies on a process initiated after the identification of a simple auditory cue -irrespective of the stimulus categorization needed for response selection-, whereas in the GNG task inhibition was applied in relation to an appropriate categorization of the stimulus.

Previous studies have pointed out methodological limitations in differentiating between inhibitory functions engaged in the GNG and SST at the brain activity level due to the existence of a high degree of functional similarity shared by these two processes (Hampshire, Chamberlain, Monti, Duncan, & Owen, 2010). Likewise, it has been postulated that although the GNG task is more related with proactive inhibition processes, whereas the SST has been described to cope with reaction inhibition, similar inhibition processes merely cooccur during behavioral inhibition, making it difficult to differentiate one from the other (Mostofsky & Simmonds, 2008).

The relationship of the inhibitory processes with food consumption has been indirectly investigated when studying the effect of inhibitory training tasks on weight loss. Two recent meta-analyses reported relevant effect sizes for the effect of inhibition training on food consumption, and interestingly, these effects were larger when using the GNG task than the SST for the training (Allom, Mullan, & Hagger, 2016; Jones et al., 2016). It has been proposed that food deprivation enhances the perceptual processing of food stimuli in later processing stages related to stimulus recognition and focused attention. In a similar vein, it has been postulated that attention is primarily driven by motivation in natural environments (Lang, Bradley, & Cuthbert, 1997). Thus, one could argue that food-related stimuli must become the focus of attention to implement appropriate behaviors to restore energy balance. Accordingly, focused attention on food cues might elicit both approach and anticipation behaviors (Seibt et al., 2007). This view has received supportive electrophysiological evidence, with modulations in early and later time windows specific for food pictures and under fasting conditions (Stockburger, Schmälzle, Flaisch, Bublatzky, & Schupp, 2009).

Since a substantial part of the nervous system in animals and humans is concerned with the procurement of food, it is possible that an effective interaction between metabolism and the brain could have evolved to engage powerful motivational drives to guarantee energy supply by creating representations and reward expectancies related to food cues (Berthoud, 2007). If so, this would explain why poorer inhibitory control has been associated with a vulnerability to food cues (Guerrieri, Nederkoorn, & Jansen, 2007; Nederkoorn, 2014). Several metabolic mediators deployed into the circulatory system to improve energy supply act first at the hypothalamus as well as the corticolimbic circuit to stimulate food procurement. Leptin has been shown to act directly on mesolimbic dopamine neurons to modulate food craving (Hommel et al., 2006). Low levels of leptin elicit a motivational drive in the hypothalamus, manifested as an increased hedonic value and an incentive salience for primary and conditioned food cues (Berthoud, 2007). Taking this idea one step further, it could be speculated that the inhibitory function is an unnecessary executive function (Hampshire et al., 2010) for controlling energy balance. In contrast, optimizing attentional monitoring and detection would be sufficient for reaching adequate proactive control of energy intake (Dehaene, Kerszberg, & Changeux, 1998; Stuss & Alexander, 2007). This hypothesis is consistent with our findings of a motivational goal evidenced in the form of an inhibitory control decline in front of food cues, and especially when they represent high-caloric items.

There is a strong impetus to fight against obesity, an eating disorder that has reached the risk of an epidemic in the last decade (James, Leach, Kalamara, & Shayeghi, 2001), as obesity has nearly tripled worldwide since 1975 (Di Cesare et al., 2016). Understanding the behavioral patterns that derive from a metabolic deficit (i.e., how metabolism and cognitive function interact with each other) is crucial to develop research plans that would lead us to create fruitful therapies or dietary plans to control and effectively restrict highly caloric food consumption, especially in populations at risk for developing morbid obesity. Taken together, the results of our study provide the first evidence that hunger is associated with some specific aspect of inhibitory processes in normal-weight participants, which might be more easily interpreted as the motivational drive to eat, initiated as a metabolic reaction to an energy deficit, might affect attentional control to food-related cues.

## Acknowledgments

This work was supported by the *Ministerio de Economía, Industria y Competitividad* (MEIC) (Grant number: PSI2016-79678) financed by the *Agencia Estatal de Investigación* (AEI) and *Fondo Europeo de Desarrollo Regional* (FEDER) from the European Union (UE).

## Data availability

The datasets generated during and/or analyzed during the current study are available from the corresponding author on request.

## References

Allom, V., Mullan, B., & Hagger, M. (2016). Does inhibitory control training improve health behaviour? A meta-analysis. Health Psychology Review, 10(2), 168–186. https://doi.org/10.1080/17437199.2015.1051078

Ballestero-Arnau, M., Moreno-Sánchez, M., & Cunillera, T. (2021). Food is special by itself: Neither valence, arousal, food appeal, nor caloric content modulate the attentional bias induced by food images. Appetite, 156. https://doi.org/10.1016/j.appet.2020.104984

Bates, D., Mächler, M., Bolker, B., & Walker, S. (2015). Fitting linear mixed-effects models using lme4. Journal of Statistical Software, 67, 1–48. https://doi.org/10.1017/CBO9781107415324.004

Baumeister, R. F. (2014). Self-regulation, ego depletion, and inhibition. Neuropsychologia, 65, 313–319. https://doi.org/10.1016/j.neuropsychologia.2014.08.012

Berthoud, H. R. (2002). Multiple neural systems controlling food intake and body weight. Neuroscience and Biobehavioral Reviews, 26(4z), 393–428. https://doi.org/10.1016/S0149-7634(02)00014-3

Berthoud, H. R. (2007). Interactions between the “cognitive” and “metabolic” brain in the control of food intake. Physiology & Behavior, 91(5), 486–498. https://doi.org/10.1016/j.physbeh.2006.12.016

Bogacz, R., Brown, E., Moehlis, J., Holmes, P., & Cohen, J. D. (2006). The physics of optimal decision making: A formal analysis of models of performance in two-alternative forced-choice tasks. Psychological Review, 113(4), 700–765. https://doi.org/10.1037/0033-295X.113.4.700

Boisseau, C. L., Thompson-Brenner, H., Caldwell-Harris, C., Pratt, E., Farchione, T., & Harrison Barlow, D. (2012). Behavioral and cognitive impulsivity in obsessive-compulsive disorder and eating disorders. Psychiatry Research, 200(2–3), 1062–1066. https://doi.org/10.1016/j.psychres.2012.06.010

Bongers, P., Van de Giessen, E., Roefs, A., Nederkoorn, C., Booij, J., Van den Brink, W., & Jansen, A. (2015). Being impulsive and obese increases susceptibility to speeded detection of high-calorie foods. Health Psychology, 34(6), 677–685. https://doi.org/10.1037/hea0000167

Castellanos, E. H., Charboneau, E., Dietrich, M. S., Park, S., Bradley, B. P., Mogg, K., & Cowan, R. L. (2009). Obese adults have visual attention bias for food cue images: Evidence for altered reward system function. International Journal of Obesity, 33(9), 1063–1073. https://doi.org/10.1038/ijo.2009.138

Claes, L., Nederkoorn, C., Vandereycken, W., Guerrieri, R., & Vertommen, H. (2006). Impulsiveness and lack of inhibitory control in eating disorders. Eating Behaviors, 7(3), 196–203. https://doi.org/10.1016/j.eatbeh.2006.05.001

Cunningham, C. A., & Egeth, H. E. (2018). The capture of attention by entirely irrelevant pictures of calorie-dense foods. Psychonomic Bulletin and Review, 25(2), 586–595. https://doi.org/10.3758/s13423-017-1375-8

Dehaene, S., Kerszberg, M., & Changeux, J. P. (1998). A neuronal model of a global workspace in effortful cognitive tasks. Proceedings of the National Academy of Sciences of the United States of America, 95(24), 14529–14534. https://doi.org/10.1073/pnas.95.24.14529

Di Cesare, M., Bentham, J., Stevens, G. A., Zhou, B., Danaei, G., Lu, Y.,… Cisneros, J. Z. (2016). Trends in adult body-mass index in 200 countries from 1975 to 2014: A pooled analysis of 1698 population-based measurement studies with 19.2 million participants. The Lancet, 387(10026), 1377–1396. https://doi.org/10.1016/S0140-6736(16)30054-X

Erdfelder, E., FAul, F., Buchner, A., & Lang, A. G. (2009). Statistical power analyses using G*Power 3.1: Tests for correlation and regression analyses. Behavior Research Methods, 41(4), 1149–1160. https://doi.org/10.3758/BRM.41.4.1149

Forestell, C. A., Lau, P., Gyurovski, I. I., Dickter, C. L., & Haque, S. S. (2012). Attentional biases to foods: The effects of caloric content and cognitive restraint. Appetite, 59(3), 748–754. https://doi.org/10.1016/j.appet.2012.07.006

Foroni, F., Pergola, G., Argiris, G., & Rumiati, R. I. (2013). The FoodCast research image database (FRIDa). Frontiers in Human Neuroscience, 7, 1–19. https://doi.org/10.3389/fnhum.2013.00051

Ganor-Moscovitz, N., Weinbach, N., Canetti, L., & Kalanthroff, E. (2018). The effect of food-related stimuli on inhibition in high vs. low restrained eaters. Appetite, 131(July), 53–58. https://doi.org/10.1016/j.appet.2018.08.037

Gerdan, G., & Kurt, M. (2020). Response inhibition according to the stimulus and food type in exogenous obesity. Appetite, 150(March), 104651. https://doi.org/10.1016/j.appet.2020.104651

Groman, S. M., James, A. S., & Jentsch, J. D. (2009). Poor response inhibition: At the nexus between substance abuse and attention deficit/hyperactivity disorder. Neuroscience and Biobehavioral Reviews, 33(5), 690–698. https://doi.org/10.1016/j.neubiorev.2008.08.008

Guerrieri, R., Nederkoorn, C., & Jansen, A. (2007). How impulsiveness and variety influence food intake in a sample of healthy women. Appetite, 48(1), 119–122. https://doi.org/10.1016/j.appet.2006.06.004

Guerrieri, R., Nederkoorn, C., Stankiewicz, K., Alberts, H., Geschwind, N., Martijn, C., & Jansen, A. (2007). The influence of trait and induced state impulsivity on food intake in normal-weight healthy women. Appetite, 49(1), 66–73. https://doi.org/10.1016/j.appet.2006.11.008

Hampshire, A., Chamberlain, S. R., Monti, M. M., Duncan, J., & Owen, A. M. (2010). The role of the right inferior frontal gyrus: inhibition and attentional control. NeuroImage, 50(3), 1313–1319. https://doi.org/10.1016/j.neuroimage.2009.12.109

Hommel, J. D., Trinko, R., Sears, R. M., Georgescu, D., Liu, Z. W., Gao, X. B.,… DiLeone, R. J. (2006). Leptin Receptor Signaling in Midbrain Dopamine Neurons Regulates Feeding. Neuron, 51(6), 801–810. https://doi.org/10.1016/j.neuron.2006.08.023

Huster, R. J., Enriquez-Geppert, S., Lavallee, C. F., Falkenstein, M., & Herrmann, C. S. (2013). Electroencephalography of response inhibition tasks: Functional networks and cognitive contributions. International Journal of Psychophysiology, 87(3), 217–233. https://doi.org/10.1016/j.ijpsycho.2012.08.001

Infection Prevention during Blood Glucose Monitoring and Insulin Administration. (n.d.). Retrieved June 27, 2019, from Centers for Disease Control and Prevention website: https://www.cdc.gov/injectionsafety/blood-glucose-monitoring.html

James, P. T., Leach, R., Kalamara, E., & Shayeghi, M. (2001). The Worldwide Obesity Epidemic. Obesity Research, 9(S11), 228–233. https://doi.org/10.1038/oby.2001.123

Jones, A., Di Lemma, L. C. G., Robinson, E., Christiansen, P., Nolan, S., Tudur-Smith, C., & Field, M. (2016). Inhibitory control training for appetitive behaviour change: A meta-analytic investigation of mechanisms of action and moderators of effectiveness. Appetite, 97, 16–28. https://doi.org/10.1016/j.appet.2015.11.013

Kirsten, H., Seib-Pfeifer, L. E., Koppehele-Gossel, J., & Gibbons, H. (2019). Food has the right of way: Evidence for prioritised processing of visual food stimuli irrespective of eating style. Appetite, 142(June), 104372. https://doi.org/10.1016/j.appet.2019.104372

Kollei, I., Rustemeier, M., Schroeder, S., Jongen, S., Herpertz, S., & Loeber, S. (2018). Cognitive control functions in individuals with obesity with and without binge-eating disorder. International Journal of Eating Disorders, 51(3), 233–240. https://doi.org/10.1002/eat.22824

Lang, P., Bradley, M., & Cuthbert, B. (1997). Motivated attention: affect, activation, and action. In R. F. Simons, P. J. Lang, & M. Balaban (Eds.), Attention and orienting: sensory and motivational processes (pp. 97–136). New York: Lawrence Erlbaum Associates, Inc.

Lavagnino, L., Arnone, D., Cao, B., Soares, J. C., & Selvaraj, S. (2016). Inhibitory control in obesity and binge eating disorder: A systematic review and meta-analysis of neurocognitive and neuroimaging studies. Neuroscience and Biobehavioral Reviews, 68, 714–726. https://doi.org/10.1016/j.neubiorev.2016.06.041

Lee, M., & Lee, J. H. (2021). Automatic attentional bias toward high-calorie food cues and body shape concerns in individuals with a high level of weight suppression: Preliminary findings. Eating Behaviors, 40(October 2018), 101471. https://doi.org/10.1016/j.eatbeh.2020.101471

Loeber, S., Grosshans, M., Herpertz, S., Kiefer, F., & Herpertz, S. C. (2013). Hunger modulates behavioral disinhibition and attention allocation to food-associated cues in normal-weight controls. Appetite, 71, 32–39. https://doi.org/10.1016/j.appet.2013.07.008

Logan, G. D., Schachar, R. J., & Tannock, R. (1997). Impulsivity and inhibitory control. Psychological Science, 8(1), 60–64. https://doi.org/10.1111/j.1467-9280.1997.tb00545.x

Lyu, Z., Zheng, P., Chen, H., & Jackson, T. (2017). Approach and inhibition responses to external food cues among average-weight women who binge eat and weight-matched controls. Appetite, 108, 367–374. https://doi.org/10.1016/j.appet.2016.10.025

Mela, D. J., Aaron, J. I., & Gatenby, S. J. (1996). Relationships of consumer characteristics and food deprivation to food purchasing behavior. Physiology and Behavior, 60(5), 1331–1335. https://doi.org/10.1016/S0031-9384(96)00241-7

Miyake, A., Friedman, N. P., Emerson, M. J., Witzki, A. H., Howerter, A., & Wager, T. D. (2000). The Unity and Diversity of Executive Functions and Their Contributions to Complex “Frontal Lobe” Tasks: A Latent Variable Analysis. Cognitive Psychology, 41(1), 49–100. https://doi.org/10.1006/cogp.1999.0734

Mogg, K., Bradley, B. P., Hyare, H., & Lee, S. (1998). Selective attention to food-related stimuli in hunger: Are attentional biases specific to emotional and psychopathological states, or are they also found in normal drive states? Behaviour Research and Therapy, 36, 227–237. https://doi.org/10.1016/S0005-7967(97)00062-4

Mostofsky, S. H., & Simmonds, D. J. (2008). Response inhibition and response selection: Two sides of the same coin. Journal of Cognitive Neuroscience, 20(5), 751–761. https://doi.org/10.1162/jocn.2008.20500

Nederkoorn, C., Guerrieri, R., Havermans, R. C., Roefs, A., & Jansen, A. (2009). The interactive effect of hunger and impulsivity on food intake and purchase in a virtual supermarket. International Journal of Obesity, 33, 905–912. https://doi.org/10.1038/ijo.2009.98

Nederkoorn, C. (2014). Effects of sales promotions, weight status, and impulsivity on purchases in a supermarket. Obesity, 22(5), E2–E5. https://doi.org/10.1002/oby.20621

Nederkoorn, C., Houben, K., Hofmann, W., Roefs, A., & Jansen, A. (2010). Control yourself or just eat what you like? weight gain over a year is predicted by an interactive effect of response inhibition and implicit preference for snack foods. Health Psychology, 29(4), 389–393. https://doi.org/10.1037/a0019921

Nederkoorn, C., Smulders, F. T. Y., Havermans, R. C., Roefs, A., & Jansen, A. (2006). Impulsivity in obese women. Appetite, 47(2), 253–256. https://doi.org/10.1016/j.appet.2006.05.008

Neimeijer, R. A. M., de Jong, P. J., & Roefs, A. (2013). Temporal attention for visual food stimuli in restrained eaters. Appetite, 64, 5–11. https://doi.org/10.1016/j.appet.2012.12.013

Nijs, I. M. T., Franken, I. H. A., & Muris, P. (2009). Enhanced processing of food-related pictures in female external eaters. Appetite, 53(3), 376–383. https://doi.org/10.1016/j.appet.2009.07.022

Nisbett, R. E., & Kanouse, D. E. (1969). Obesity, food deprivation, and supermarket shopping behavior. Journal of Personality and Social Psychology, 12(4), 289–294. https://doi.org/10.1037/h0027799

Oliva, R., Morys, F., Horstmann, A., Castiello, U., & Begliomini, C. (2019). The impulsive brain: Neural underpinnings of binge eating behavior in normal-weight adults. Appetite, 136, 33–49. https://doi.org/10.1016/j.appet.2018.12.043

Ratcliff, R. (1993). Methods for dealing with response time outliers. Psychological Bulletin, 114(3), 510–532.

Seibt, B., Häfner, M., & Deutsch, R. (2007). Prepared to eat: How immediate affective and motivational responses to food cues are influenced by food deprivation. European Journal of Social Psychology, 37(2), 359–379. https://doi.org/10.1002/ejsp.365

Stockburger, J., Schmälzle, R., Flaisch, T., Bublatzky, F., & Schupp, H. T. (2009). The impact of hunger on food cue processing: An event-related brain potential study. NeuroImage, 47, 1819–1829. https://doi.org/10.1016/j.neuroimage.2009.04.071

Stuss, D. T., & Alexander, M. P. (2007). Is there a dysexecutive syndrome? Philosophical Transactions of the Royal Society B: Biological Sciences, 362(1481), 901–915. https://doi.org/10.1098/rstb.2007.2096

Swinburn, B. A., Sacks, G., Hall, K. D., McPherson, K., Finegood, D. T., Moodie, M. L., & Gortmaker, S. L. (2011). The global obesity pandemic: Shaped by global drivers and local environments. The Lancet, 378(9793), 804–814. https://doi.org/10.1016/S0140-6736(11)60813-1

Van Dillen, L. F., Papies, E. K., & Hofmann, W. (2013). Turning a blind eye to temptation: How cognitive load can facilitate self-regulation. Journal of Personality and Social Psychology, 104(3), 427–443. https://doi.org/10.1037/a0031262

Verbruggen, F., Aron, A. R., Band, G. P. H., Beste, C., Bissett, P. G., Brockett, A. T.,… Boehler, C. N. (2019). A consensus guide to capturing the ability to inhibit actions and impulsive behaviors in the stop-signal task. ELife, 8, 1–26. https://doi.org/10.7554/eLife.46323

Verbruggen, F., & Logan, G. D. (2009). Models of response inhibition in the stop-signal and stop-change paradigms. Neuroscience and Biobehavioral Reviews, 33(5), 647–661. https://doi.org/10.1016/j.neubiorev.2008.08.014

Verbruggen, F., & Logan, G. D. (2017). Control in Response Inhibition. In Wiley Online Books. The Wiley Handbook of Cognitive Control (pp. 97–110). https://doi.org/10.1002/9781118920497.ch6

Verbruggen, F., McLaren, I. P. L., & Chambers, C. D. (2014). Banishing the Control Homunculi in Studies of Action Control and Behavior Change. Perspectives on Psychological Science, 9(5), 497–524. https://doi.org/10.1177/1745691614526414

Weinbach, N., Lock, J., & Bohon, C. (2020). Superior response inhibition to high-calorie foods in adolescents with anorexia nervosa. Behaviour Research and Therapy, 124(July 2019), 103441. https://doi.org/10.1016/j.brat.2019.103441

Werthmann, J., Roefs, A., Nederkoorn, C., Mogg, K., Bradley, B. P., & Jansen, A. (2011). Can(not) Take my Eyes off it: Attention Bias for Food in Overweight Participants. Health Psychology, 30(5), 561–569. https://doi.org/10.1037/a0024291

WHO. (2016). Obesity and overweight: Fact sheet 311. https://doi.org/Retrieved from http://www.who.int/mediacentre/factsheets/fs311/en/

Wonderlich, S. A., Connolly, K. M., & Stice, E. (2004). Impulsivity as a risk factor for eating disorder behavior: Assessment implications with adolescents. International Journal of Eating Disorders, 36(2), 172–182. https://doi.org/10.1002/eat.20033

Wu, M., Hartmann, M., Skunde, M., Herzog, W., & Friederich, H. C. (2013). Inhibitory control in bulimic-type eating disorders: A systematic review and meta-analysis. PLoS ONE, 8(12). https://doi.org/10.1371/journal.pone.0083412

Zeltser, L. M., Seeley, R. J., & Tschöp, M. H. (2012). Synaptic plasticity in neuronal circuits regulating energy balance. Nature Neuroscience, 15, 336–1342. https://doi.org/10.1038/nn.3219

Zheng, H., & Berthoud, H.-R. (2008). Neural Systems Controlling the Drive to Eat: Mind Versus Metabolism. Physiology, 23(2), 75–83. https://doi.org/10.1152/physiol.00047.2007

